# scnanoseq: an nf-core pipeline for Oxford Nanopore single-cell RNA-sequencing

**DOI:** 10.1101/2025.04.08.647887

**Authors:** Austyn Trull, nf-core community, Elizabeth A. Worthey, Lara Ianov

## Abstract

Recent advancements in long-read single-cell RNA sequencing (scRNA-seq) have facilitated the quantification of full-length transcripts and isoforms at the single-cell level. Historically, long-read data would need to be complemented with short-read single-cell data in order to overcome the higher sequencing errors to correctly identify cellular barcodes and unique molecular identifiers. Improvements in Oxford Nanopore sequencing, and development of novel computational methods have removed this requirement. Though these methods now exist, the limited availability of modular and portable workflows remains a challenge. Here we present, nf-core/scnanoseq, a secondary analysis pipeline for long-read single-cell and single-nuclei RNA that delivers gene and transcript-level quantification. The scnanoseq pipeline is implemented using Nextflow and is built upon the nf-core framework, enabling portability across computational environments, scalability and reproducibility of results across pipeline runs. The nf-core/scnanoseq workflow follows best practices for analyzing single-cell and single-nuclei data, performing barcode detection and correction, genome and transcriptome read alignment, unique molecular identifier deduplication, gene and transcript quantification, and extensive quality control reporting.

**AVAILABILITY OF DATA AND MATERIALS:** nf-core/scnanoseq is available at https://github.com/nf-core/scnanoseq under the MIT License and the documentation is available at https://nf-co.re/scnanoseq. The downstream analytical code for validation analysis is available at https://github.com/U-BDS/scnanoseq_analysis and all dataset sources have been disclosed under the methods section for each respective dataset.

## INTRODUCTION

Advances in single-cell and single-nuclei transcriptomics have enhanced our understanding of cellular heterogeneity by enabling high-resolution gene expression analysis. Traditionally, single cell RNA sequencing (scRNA-seq) relied on short-read sequencing, which provided high base accuracy but failed to capture full-length transcripts (Byrne *et al*. 2017, Gupta *et al*. 2018, Tian *et al*. 2021, Shi *et al*. 2023). Long-read platforms from Pacific Biosciences and Oxford Nanopore Technologies (ONT) were available and capable of full-length transcript sequencing, but were limited by higher error rates and lower throughput, which complicated barcode and Unique Molecular Identifier (UMI) recovery. Recent improvements, such as ONT’s Q20+ chemistry (10X Genomics 2022c), have, overcome these limitations, improving accuracy and enabling isoform-level quantification without the need for complementary short-read sequencing (10X Genomics 2022c, Prjibelski *et al*. 2023, Shi *et al*. 2023, Kumari *et al*. 2024).

Computational workflows, such as FLAMES (Tian *et al*. 2021), wf-single-cell (Oxford Nanopore Technologies 2024), and scywalker (De Rijk *et al*. 2024), support long-read-based single-cell quantification but often rely on custom tooling or *ad hoc* workflows, hindering reproducibility, limiting configurability and reporting, and complicating integration with broader bioinformatics workflows. Adapting to new data, parameters, or references can be slow, and inconsistent reporting and poor modularity can limit scalability and result comparison. To overcome these challenges, we developed nf-core/scnanoseq, a Nextflow (Di Tommaso *et al*. 2017)-based pipeline within the nf-core framework (Ewels *et al*. 2020, Langer *et al*. 2024) for single-cell ONT data. It integrates open-source tools for gene- and transcript-level quantification without short-read dependency and offers genome- and transcriptome-aligned analysis. Built with Nextflow DSL 2.0 best practices, the pipeline is modular, configurable, and portable, allowing users to tailor workflows while maintaining reproducibility. These features make nf-core/scnanoseq a robust and user-friendly solution for long-read single-cell RNA sequencing.

### PIPELINE DESIGN AND IMPLEMENTATION

nf-core/scnanoseq is built with Nextflow (Di Tommaso *et al*. 2017) DSL 2.0, ensuring modularity and simplifying future expansions. It leverages the nf-core (Ewels *et al*. 2020, Langer *et al*. 2024) framework, providing standardized guidelines and essential tools, including automated testing. The pipeline runs seamlessly on local machines, High Performance Computers (HPC), and cloud environments, with built-in support for Docker and Singularity, ensuring reproducibility and portability without manual software installation. Designed as an end-to-end solution, nf-core/scnanoseq processes ONT 10X Genomics single-cell/nuclei data **(Fig. 1)**. The following sections detail its components.

**Figure 1:**
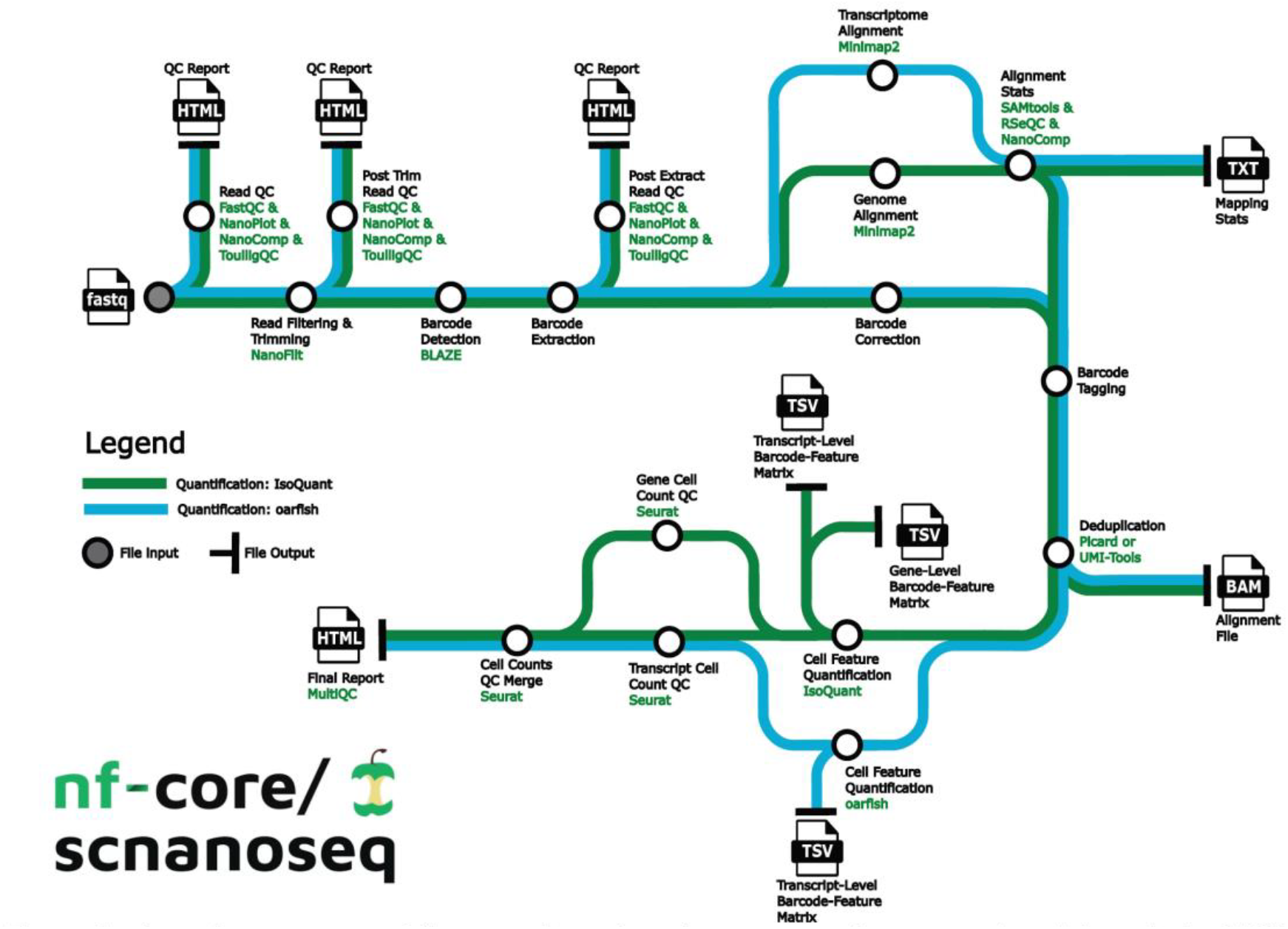
nf-core/scnanoseq workflow overview. nf-core/scnanoseq performs secondary data analysis of 10X Genomics single-celUnude data derived from Oxford Nanopore sequencing. The diagram outlines the pathways to gene-level and transcript-level analysis, with file outputs noted by the file type

### Initial FASTQ Processing

nf-core/scnanoseq accepts FASTQ files and sample metadata as input, requiring reads to contain a cellular barcode and UMI. While currently validated for use with 10X Genomics sequencing kits, its modular design allows for future support of other barcode formats. Optionally, FASTQ files can be trimmed and filtered using NanoFilt (De Coster *et al*. 2018) based on read quality and length. The trimmed or unprocessed FASTQ files are then processed with BLAZE (You *et al*. 2023) to extract uncorrected cellular barcodes and UMIs.

### Barcode Processing and Alignment

Barcodes, UMIs, and additional sequences (e.g., PCR primers, TSO, poly-T tails) are extracted from reads using a custom script, leveraging raw BLAZE (You *et al*. 2023) outputs. This step generates a barcode-free FASTQ file for alignment with Minimap2 (Li 2018) and a CSV containing barcodes, UMIs, and quality scores for correction.

Barcode correction runs in parallel with alignment using a custom Python script. Barcodes are ranked by abundance and corrected if they fall within a Hamming distance of ≤2 from a whitelist barcode and have a posterior probability ≥97.5%. Corrected barcodes are updated accordingly.

After alignment and correction, a custom script tags each BAM read with raw and corrected barcodes (CR, CB), UMI (UR), and quality scores (CY, UY). The BAM file is deduplicated using UMI-Tools (Smith *et al*. 2017) or Picard MarkDuplicates (Broad 2019). To optimize UMI-Tools, genome-aligned BAMs are split by chromosome, while transcriptome-aligned BAMs are split by features belonging to the same chromosome. Deduplication is optional when using IsoQuant (Prjibelski *et al*. 2023) but required for oarfish (Jousheghani and Patro 2024).

### Quantification

The barcode and UMI-tagged BAM files are then input for nf-core/scnanoseq quantification. The pipeline supports two methods, which can be run individually or in parallel. IsoQuant (Prjibelski *et al*. 2023) performs gene- and transcript-level quantification using genome-aligned BAMs. The ‘--read_groups’ flag groups results by cellular barcode, generating barcode-feature matrices for downstream analysis with tertiary analysis packages, such as Seurat (Hao *et al*. 2021) or Scanpy (Wolf *et al*. 2018). Oarfish (Jousheghani and Patro 2024) quantifies transcripts from transcriptome-aligned, UMI-deduplicated BAMs. Its outputs are further processed to produce results compatible with single-cell analysis tools, including those within this pipeline.

### Quality Control

nf-core/scnanoseq performs quality control (QC) at multiple stages: raw data, post-mapping, and post-quantification. For FASTQ files, FastQC (Andrews 2010), NanoPlot (De Coster and Rademakers 2023), NanoComp (De Coster and Rademakers 2023), and ToulligQC (Dias *et al*. 2024) provide general and long-read-specific QC across three FASTQ sets: (1) raw input, (2) trimmed, and (3) barcode-extracted. Reports and QC images are generated for review. For BAM files, SAMtools (Li *et al*. 2009), RSeQC (Wang *et al*. 2012), and NanoComp provide post-mapping QC. SAMtools flagstat, idxstats, and stats are run on (1) initially mapped BAMs, (2) barcode- and UMI-tagged BAMs, and (3) UMI-deduplicated BAMs.

Custom QC steps further extend these analyses. After quantification, Seurat (Hao *et al*. 2021) calculates single-cell metrics (e.g., cell counts, mean reads per cell, nFeature/nCount plots). A summary CSV tracks read counts across key steps. Finally, MultiQC (Ewels *et al*. 2016) compiles QC results, including custom metrics, into a single HTML report, for streamlined review.

## BENCHMARK AND VALIDATION

nf-core/scnanoseq has been extensively benchmarked and validated as shown in the <u>**Supplementary Information**</u>.

## CONCLUSION

Here, we present nf-core/scnanoseq, a robust pipeline for long-read single-cell RNA-seq analysis that enables both gene- and isoform-level transcriptomic profiling through genome- and transcriptome-based quantification workflows. Fully automated, nf-core/scnanoseq streamlines preprocessing steps including alignment, deduplication, barcode correction, and tagging, while ensuring reproducibility, portability and scalability.

Unlike other long-read single-cell pipelines such as scywalker (De Rijk *et al*. 2024) and wf-single-cell (Oxford Nanopore Technologies 2024), nf-core/scnanoseq leverages the nf-core framework to deliver modular, peer-reviewed workflows supporting flexible and reproducible analysis. The use of Nextflow DSL 2.0 enables seamless expansion, allowing customization of key steps, including FASTQ trimming, filtering, and barcode whitelists. Furthermore, nf-core/scnanoseq uniquely supports two quantification methods, IsoQuant (Prjibelski *et al*. 2023) and oarfish (Jousheghani and Patro 2024), within a unified framework, minimizing manual intervention while offering flexibility based on their specific needs.

Ongoing and planned improvements include updating supported tools like IsoQuant and BLAZE (You *et al*. 2023) as new versions are released, and optimizations to improve pipeline efficiency, particularly for IsoQuant. Emerging methods, for example lr-kallisto (Loving *et al*. 2025) for single-cell expression quantification, will be regularly reviewed for potential integration. By combining standardized secondary analysis practices with a modular, customizable design, nf-core/scnanoseq enables accurate gene and transcript quantification while offering users the flexibility to adapt the pipeline to their specific analysis needs.

## ACKNOWLEDGMENTS

This work was supported by 3P30CA013148-48S8, Dr. Worthey start-up funds from UAB, and L.I. was supported by the Civitan International Research Center. We also acknowledge support from the University of Alabama at Birmingham Biological Data Science Core, RRID:SCR_021766. The authors also acknowledge the support from the nf-core community for developing and maintaining the nf-core infrastructure. The authors gratefully acknowledge the resources provided by the University of Alabama at Birmingham IT-Research Computing group for high performance computing (HPC) support and CPU time on the Cheaha compute cluster. This work was supported in part by the National Science Foundation under Grants Nos. OAC-1541310, the University of Alabama at Birmingham, and the Alabama Innovation Fund. Any opinions, findings, and conclusions or recommendations expressed in this material are those of the authors and do not necessarily reflect the views of the National Science Foundation or the University of Alabama at Birmingham.

## AUTHOR CONTRIBUTIONS

L.I. and E.A.W. obtained funding. A.T and L.I. designed and developed nf-core/scnanoseq. A.T. and L.I. performed validation, benchmarking and data visualization as detailed. A.T. and L.I drafted the manuscript. L.I. supervised all work. E.A.W provided feedback and revised the manuscript. All authors approved of the final manuscript.

## SUPPLEMENTARY INFORMATION

### Benchmark

Ensuring an efficient end-to-end runtime is a key feature of computational pipelines. Individual pipeline steps were benchmarked using the 10X 3’ PBMC dataset to assess time and memory efficiency (**Supplementary Figure 1A**). NanoComp (De Coster and Rademakers 2023) and SAMtools (Li *et al*. 2009) were excluded from visualization. NanoComp was omitted since it aggregates results across multiple samples, leading to high memory usage in large-scale analyses without adding unique insights for single samples. While this QC step can provide additional summary statistics, its omission does not compromise essential quality control metrics, as nf-core/scnanoseq integrates key QC results into MultiQC (Ewels *et al*. 2016) alongside other QC-specific steps. SAMtools was excluded due to the number of times various subtools are called, each contributing minimally to overall runtime.

Among QC processes, pre-alignment QC steps such as FastQC (Andrews 2010), NanoPlot (De Coster and Rademakers 2023), ToulligQC (Dias *et al*. 2024) require the most time, with FastQC being the most time-consuming. To optimize execution time, all QC steps in nf-core/scnanoseq are optional, allowing users to skip individual steps or disable all QC with a single flag. This flexibility enables users to select specific QC tools instead of running all available options, such as choosing between FastQC or NanoPlot for read QC metrics reported in MultiQC. Steps marked with an asterisk (* in Supplementary Fig. 1) indicate processes that have been parallelized, with the longest subprocess runtime reported. Parallelization in nf-core/scnanoseq is implemented through file-splitting, allowing tools to process smaller chunks of data concurrently. This approach is necessary for methods that lack native multithreading support, such as NanoFilt (De Coster *et al*. 2018), as well as for multithreaded tools that still require significant runtime, such as IsoQuant (Prjibelski *et al*. 2023).

To further evaluate the impact of Nextflow (Di Tommaso *et al*. 2017)-based file-splitting parallelization, analysis runtimes were assessed using the Shiau, CK., Lu, L., Kieser, R. et al. dataset (Shiau *et al*. 2023)(**Supplementary Fig. 1B**). The results indicate that most split processes complete within an hour, with the exception of IsoQuant, which has a maximum runtime of five hours. However, IsoQuant runtime is *heavily dependent on dataset depth*, and samples with greater sequencing depth than those presented in this work are expected to require longer runtimes. The results demonstrate that file-splitting parallelization effectively minimizes processing time across multiple steps in the pipeline. Additionally, users can further reduce runtime by selecting either IsoQuant or oarfish (Jousheghani and Patro 2024) for quantification rather than running both, allowing nf-core/scnanoseq to execute a single pathway instead of both quantifiers. The reported times in **Supplementary Figure 1** represent pipeline execution runtimes and do not account for queue times on shared HPC systems, which are subject to external factors such as overall system load and compute resource availability.

**Supplementary Figure 1:**
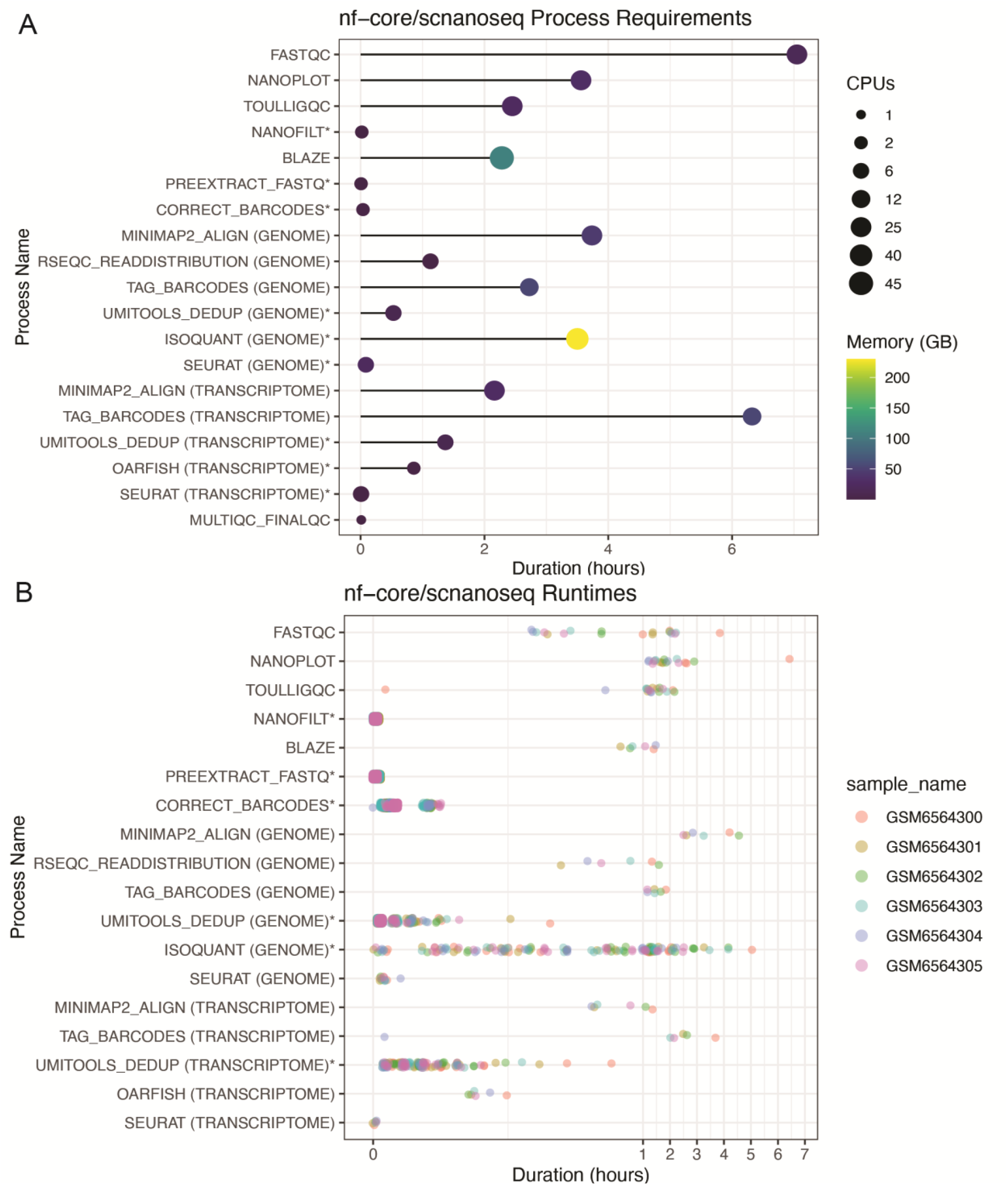
nf-core/scnanoseq benchmarks. Computational benchmark of processes associated with data processing for the genome and transcriptome paths, various QC steps, and reporting for the **A**. 3’ PBMC dataset (129.3M reads) **B**. Shiau et al dataset (105.3M-67.4M reads) Processes with an have been parallerized by nf-core/scnanoseq. where the longest process is shown in A vzhile all are shown in B. The executation time in hours is displayed in the x-axis

### nf-core/scnanoseq enables reliable quantification of long-read single-cell and nuclei datasets

nf-core/scnanoseq was evaluated across three datasets: 1. 10X Genomics 3’ PBMC dataset (10X Genomics 2022c, 10X Genomics 2022b) 2. 10X Genomics 5’ stage III squamous cell lung carcinoma (10X Genomics 2022c, 10X Genomics 2022a)(lung cancer DTCs) 3. You et al. pluripotent stem cells undergoing cortical neuronal differentiation (You *et al*. 2023) (**Supplementary Table 1**). The 3’ PMBC and 5’ lung cancer DTCs datasets included matched Illumina data for gene-level comparisons, while You et al. provided gene- and transcript-level quantification via BLAZE (You *et al*. 2023)-FLAMES (Tian *et al*. 2021) enabling direct comparisons at both levels.

In the 3’ PBMC and 5’ lung cancer DTCs datasets, Azimuth (Hao *et al*. 2021) cell annotation post-integration showed high concordance between nf-core/scnanoseq and Cell Ranger (CR) Illumina short-read data (**Supplementary Fig. 2A, 3A**), with successful cell label transfer to transcript-level data (**Supplementary Fig. 2B, 3B**). A direct comparison of detected features per cell yielded a strong correlation (Pearson r = 0.98) between nf-core/scnanoseq and CR gene-level datasets (**Supplementary Fig. 2C, 3C**). Expression analysis of Azimuth markers at both gene and transcript levels revealed comparable expression profiles between nf-core/scnanoseq and CR (**Supplementary Fig. 2D, 3D**) and isoform-specific patterns across available quantifiers (**Supplementary Fig. 4A-B**). For instance, while *CD79A* is broadly expressed in B cells, *CD79A*.*201* is the predominant isoform (**Supplementary Fig. 2D-E**). Similarly, *C1QB* is widely expressed in macrophages with *C1QB*.*201* and *C1QB*.*203* being the dominant isoforms over *C1QB*.*202* and *C1QB*.*204* (**Supplementary Fig. 3D-E**).

Comparison with You et al. dataset (BLAZE-FLAMES) showed that nf-core/scnanoseq reproduced cell clustering at both gene and transcript levels (**Supplementary Fig. 5A**) while recovering 2-3 times more features and molecules per cell in the PromethION sample (ERR9958135) (**Supplementary Fig. 5B-D**). Additionally, feature and molecule count per cell remained highly correlated at both levels (Pearson r = 0.94-0.95, **Supplementary Fig. 5C-D**).

**Supplementary Figure 2:**
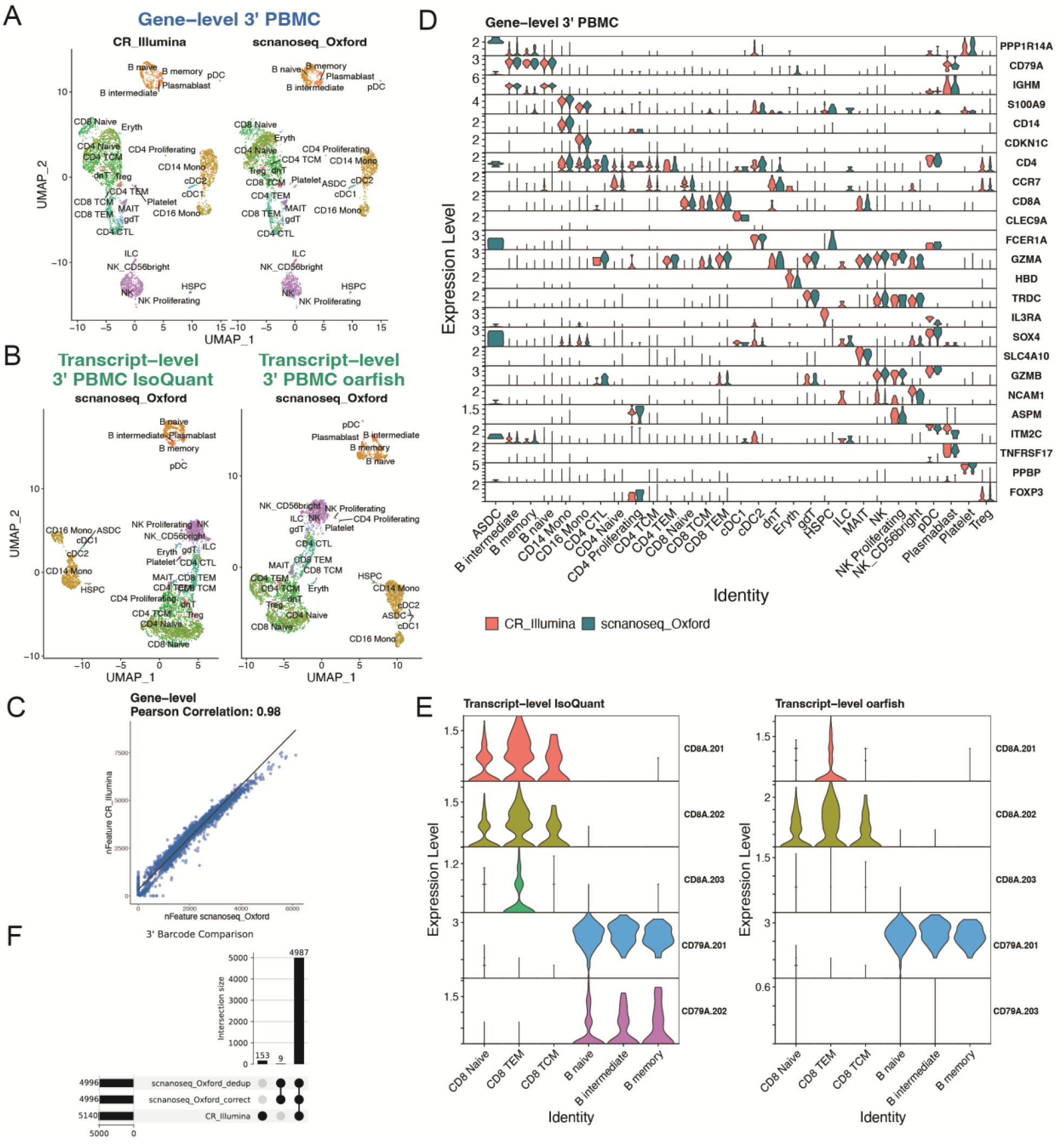
nf-core/scnanoseq validation against 3’ PBMC ground truth. UMAPs of cell dusters identified across scnanoseq and Cell Ranger [CR] short-read data at the **A**. gene-level (blue, CR and scnanoseq pipelines) and **B**. trancript-level (green, scnanoseq pipeline only, under IsoQuant and oarfish quantifiers). **C**. Gene-level scatter plot with Pearson correlation value between the number of features detected in a cell from scnanoseq (x-axis) and CR (y-axis) **D**. Gene-level expression of subset of Azimuth PBMC markers split by scnanoseq with tong-read data and CR short-read data. **E**. Violin plots comparing transcnpt-level soiorm expression for *CD8A* and *CD79A* isoforms with scnanoseq using the IsoQuant and oarfish quantifiers **F**. Barcode comparison from scnanoseq at two pipelines stages (correction and post deduplication) and CR data. Bar chart at the left represents the total number of barcodes across each dataset Bar chart at the top represents the intersection

**Supplementary Figure 3:**
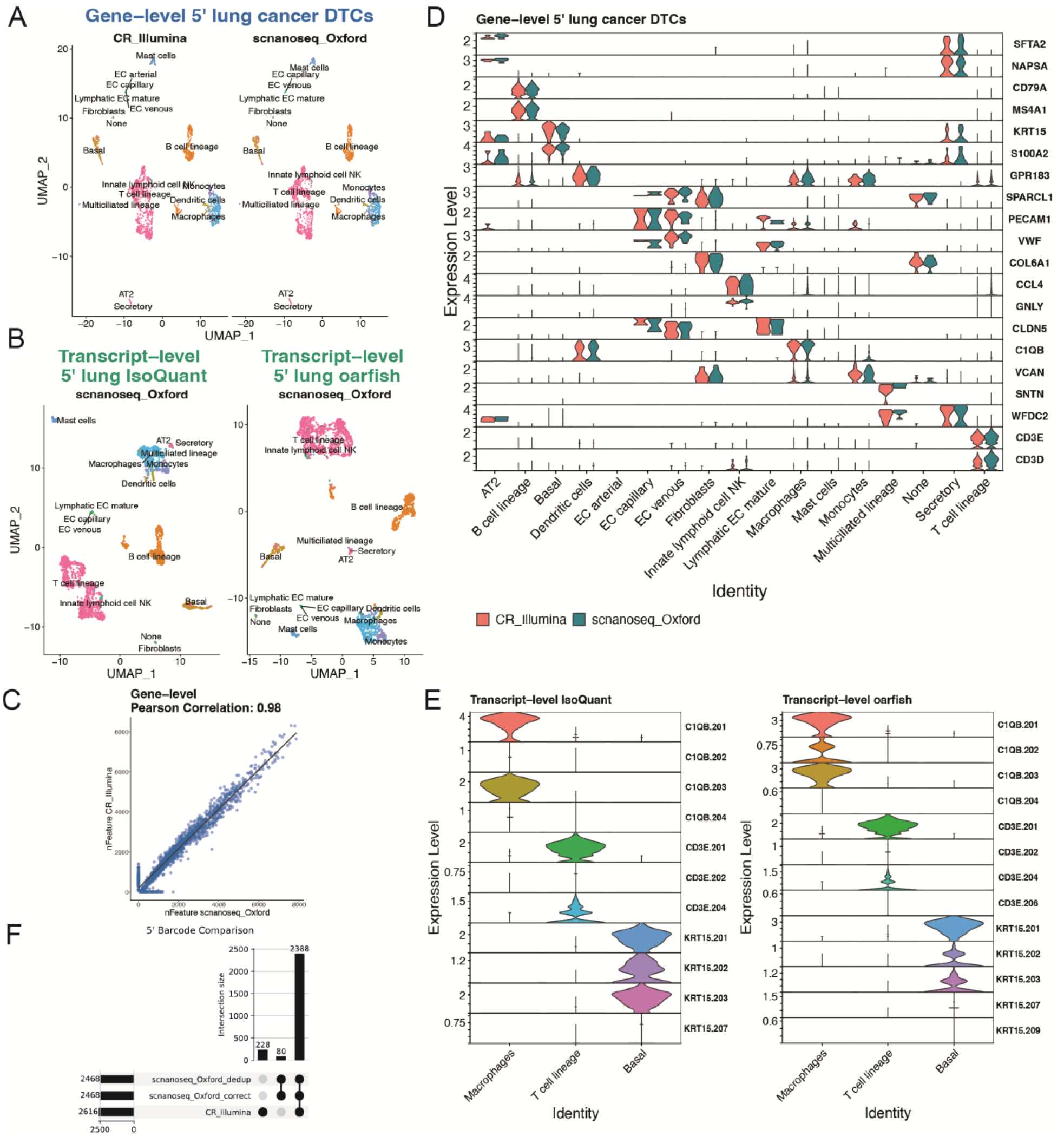
nf-core/scnanoseq validation against 5’ lung cancer DTCs ground truth. UMAPs of cell clusters identified across scnanoseq and Cell Ranger [CR] short-read data at the **A**. gene-level (blue. CR and scnanoseq pipelines) and **B**. trancript-level (green, scnanoseq pipeline only, under IsoQuant and oarfish quantifiers). **C**. Gene-level scatter plot with Pearson correlation value between the number of features detected in a cell from scnanoseq (x-axis) and CR(y-axis). **D**. Gene-level expression of subset of Azimuth lung markers split by scnanoseq with long-read data and CR short-read data. **E**. Violin plots comparing transcript-level isoform expression for *C1QB* and *CD3E* isoforms with scnanoseq using the IsoQuant and oarfish quantifiers. **F**. Barcode comparison from scnanoseq at two pipelines stages (correction and post deduplication) and CR data. Bar chart at the left represents the total number of barcodes across each dataset. Bar chart at the top represents the intersection.

**Supplementary Figure 4:**
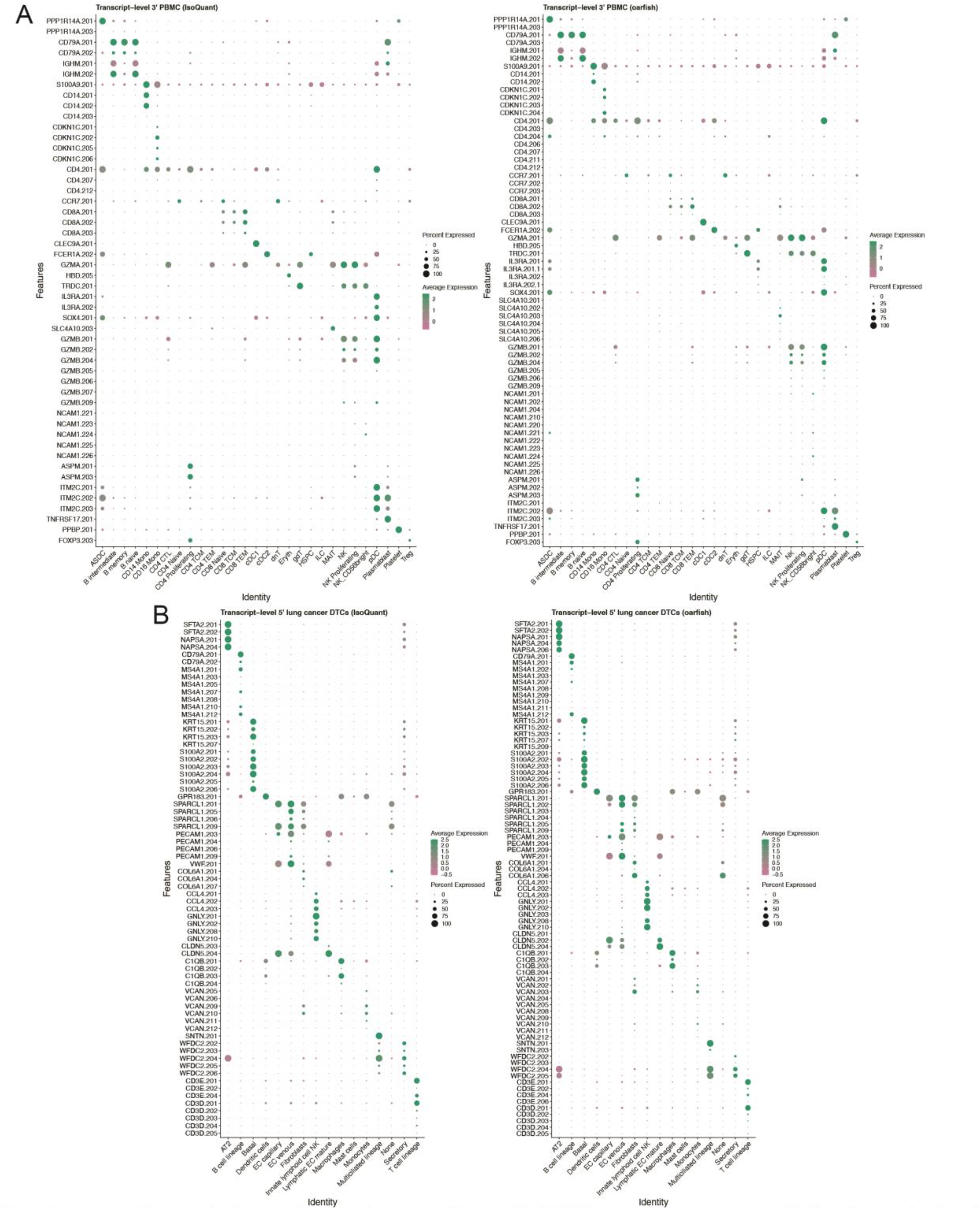
Transcript-level expression in 3’ PBMC and 5’ lung cancer DTCs. Transcript-level expression of a subset of Azimuth PBMC **(A)** and lung **(B)** markers from scnanoseq with both quantifiers processing enabled. Transcripts shown match genes displayed in Supplementary Figures 2 and 3.

**Supplementary Figure 5:**
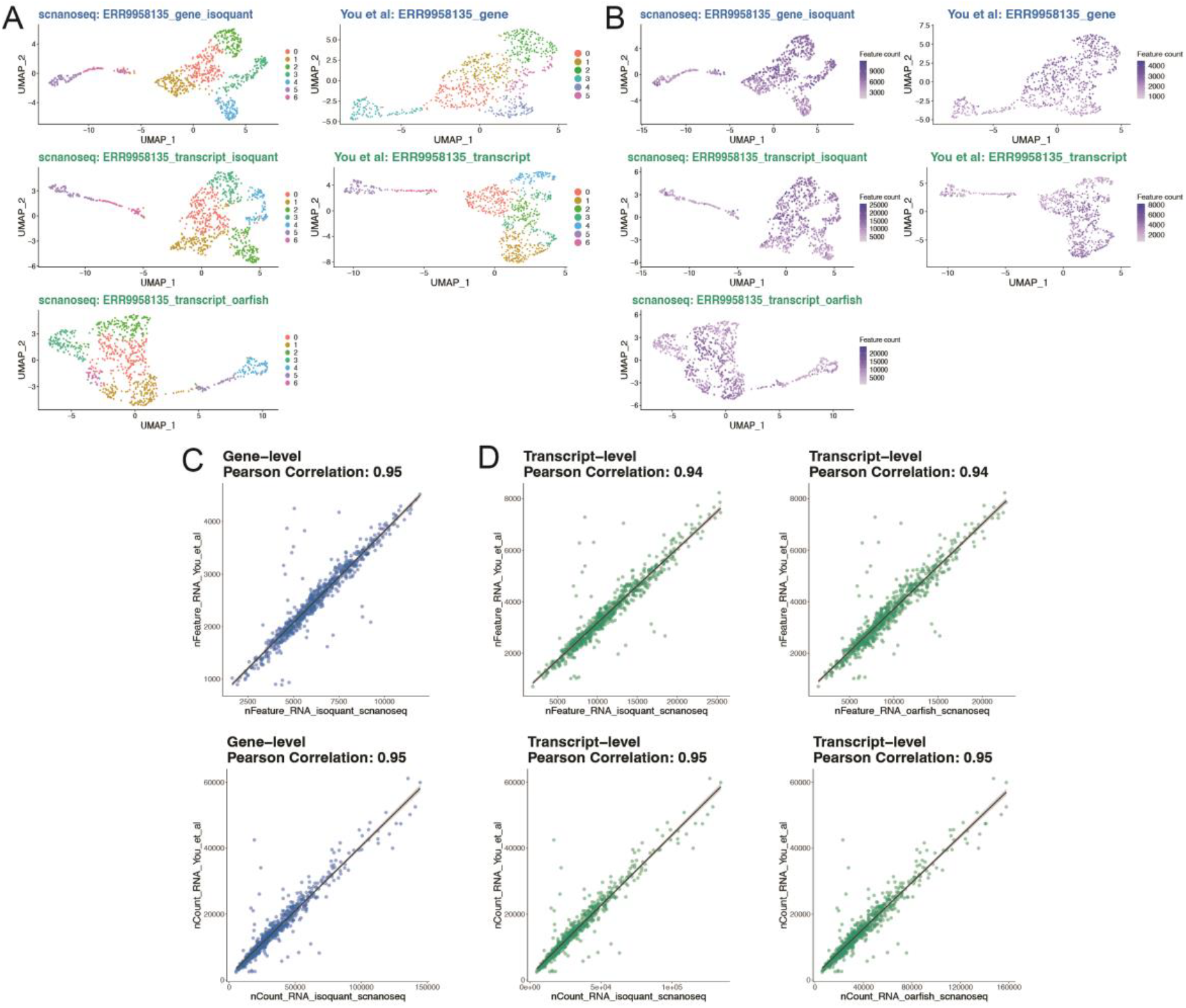
nf-core/scnanoseq re-analysis of You et al. cortical neuron culture. Sample to sample comparison (PromethlON sample ERR9958135) from scnanoseq vs. You et al. FLAMES pipeline outputs. **A**. Gene and transcript-level UMAPs of cell clusters identified across scnanoseq (top) and You et al. (bottom). **B**. Number of features detected per cell in each dataset. Each plot is scaled by dataset. **C-D**. Scatter plots with Pearson correlation values between matched cells from scnanoseq (x-axis) and You et al. (y-axis) for the number of features (nFeature) and the number of molecules (nCount) at the gene **(C)** and transcript **(D)** level.

**Supplementary Table 1:** Compiled sample name, platform, tissue type, data source and read count for all publicly available datasets used for benchmarking and validation of nf-core/scnanoseq.

| sample_name | platform | tissue_type | source | read_count | Analysis type |
| --- | --- | --- | --- | --- | --- |
| ERR9958135 | PromethION | cortical neuronal differentiated stem cells | You et al. | 61,967,455 | validation |
| 5' lung cancer DTCs | PromethION | lung cancer DTCs | 10X Application Note | 106,105,266 | validation and benchmarking |
| 3' PBMC | PromethION | PBMC | 10X Application Note | 129,264,682 | validation and benchmarking |
| GSM656300 | PromethION | A375 Cell Line | Shiau et al. | 105,347,205 | benchmarking |
| GSM656301 | PromethION | H2030 Cell Line | Shiau et al. | 67,436,356 | benchmarking |
| GSM656302 | PromethION | Primary RCC Tumor | Shiau et al. | 98,349,656 | benchmarking |
| GSM656303 | PromethION | RCC brain metastasis kidney tumor | Shiau et al. | 87,800,959 | benchmarking |
| GSM656304 | PromethION | Primary melanoma | Shiau et al. | 72,828,041 | benchmarking |
| GSM656305 | PromethION | Melanoma Brain metastasis | Shiau et al. | 98,363,541 | benchmarking |

## METHODS

### Input Datasets

To validate and benchmark nf-core/scnanoseq, we tested it on four datasets (**Supplementary Table 1):** 10X Genomics 3’ PBMC (10X Genomics 2022c, 10X Genomics 2022b) and the 5’ lung cancer DTCs (10X Genomics 2022c, 10X Genomics 2022a) as well as FASTQ files from You et al. (You *et al*. 2023) (ENA project PRJEB54718 (Leinonen *et al*. 2011)), and Shiau et al. (Edgar *et al*. 2002) (GEO accession GSE212945 (Shiau *et al*. 2023)). The latter two datasets were retrieved using nf-core/fetchngs v. 1.10.0 (Patel *et al*. 2023).

### Reference Genome, Transcriptome and Annotation

The reference genome, transcriptome, and annotation (GTF) files used for validation matched those from the original studies: GRCh38 GENCODE release 32 (Ensembl 98) (Frankish *et al*. 2022) for the 3’ PBMC, 5’ lung cancer DTC, and Shiau et al. datasets, and GRCh38 GENCODE release 31 (Ensembl 97) for the You et al. dataset.

### nf-core/scnanoseq

The nf-core/scnanoseq version 1.1.0 was used to analyze all datasets with default parameters, except for ‘min_length’ (set to 500 for all runs) and ‘split_amount’ (set to 500000 for the 3’ PBMC and 5’ lung cancer DTCs, and 1000000 for the Shiau et al. datasets). Parameters were adjusted to match the specified genome and transcriptome references, as well as the expected barcode format (3’ or 5’). For the Shiau et al. dataset, a custom ‘whitelist’ (“737K-arc-v1.txt” from 10X Genomics) was used, in line with the authors’ library preparation protocol. All runs were executed on the University of Alabama at Birmingham HPC cluster.

### Validation analysis

All post-secondary analysis results (e.g., nf-core/scnanoseq outputs or author-reported data) were processed in a Singularity container containing downstream analytical dependencies. Analysis code is available at: https://github.com/U-BDS/scnanoseq_analysis. The scope of the validation analysis focused on standard downstream procedures, such as quality control, filtering, integration (where applicable), dimensionality reduction, clustering, annotation, and visualization across matched datasets. Additional analyses, such as doublet identification, were excluded to minimize downstream transformations. Below is the validation analysis for each dataset:

<u>3’ PBMC, 5’ lung cancer DTCs:</u> The input data included nf-core/scnanoseq outputs (gene- and transcript-level matrices from both quantifiers) and sample-matched Illumina short-read data from 10X Genomics (Cell Ranger v. 7.0.1). All analyses were conducted with Seurat (v. 4.4.0) (Hao *et al*. 2021) in R (v. 4.3.2). Quality control removed low-quality cells based on criteria: nFeature > 500 (gene), nFeature > 800 (transcript), nCount > 200 (gene, transcript), percent mitochondrial < 10 (gene, transcript). Data was normalized with SCTransform V2 (‘vars.to.regress’ set to percent mitochondrial), and integration was performed with Harmony (v. 1.2.0) (Korsunsky *et al*. 2019) for gene-level data (nf-core/scnanoseq and CR_Illumina). Transcript-level data from nf-core/scnanoseq was processed separately. Additionally, Azimuth (v. 0.4.6) PBMC (Stuart *et al*. 2019) and lung (Sikkema *et al*. 2023) reference sets were implemented to annotate the cell types and the cell annotations from the gene-level dataset were transferred to the transcript-level data based on matching barcodes. Dimensionality reduction and clustering were executed with the following parameters: dims = 25, resolution = 0.7 (gene, PBMC), dims = 27, resolution = 0.6 (gene, lung), dims = 25, resolution = 0.8 (transcript, PBMC), dims = 25, resolution = 0.6 (transcript, lung). The Azimuth cell annotation was set to the predicted “celltype.I2” (PBMC) and “ann_level_3” (lung). The gene-level nFeature scatter plots were generated between all cells detected in the input data (pre-quality filtering) and the correlation values were computed with the Pearson correlation coefficient. A representative subset of Azimuth PBMC and lung markers were selected for visualization across nf-core/scnanoseq (gene, transcript) and CR_Illumina (gene).

UpSet plots were generated with the upsetplot package (v. 0.9.0) in Python (v. 3.12.0), using unique barcodes from the barcode-corrected and UMI-deduplicated BAM files. SAMtools (v.1.12) was used to extract tagged barcodes from the BAM outputs of nf-core/scnanoseq.

<u>You et al.</u>: The input data contained the outputs of nf-core/scnanoseq (gene and transcript-level matrices) and the author provided matrices (gene and transcript-level matrices). Each pipeline dataset was separately processed with quality and control filtering low-quality cells based on nFeature > 100 (gene, nf-core/scnanoseq and You et al.), nFeature > 700 (transcript, nf-core/scnanoseq and You et al.), percent mitochondrial < 20 (gene, transcript, nf-core/scnanoseq only). No percent mitochondrial filtering was performed in the You et al. data provided by the authors due to the lack of matches between known mitochondrial gene and transcript IDs in this dataset. The data was normalized with SCTransform V2 and dimensionality reduction and clustering were executed with the following parameters: dims = 6, resolution = 0.6 (gene, transcript, nf-core/scnanoseq), dims = 6, resolution = 0.7 (gene, transcript, You et al.). Data visualization and reporting were focused on the PromethION sample (ERR9958135), with nFeature and nCount scatter plots and Pearson correlation coefficient calculations.

